# Genomic characterization of a unique population of *Leishmania tropica* in a new emerging focus of cutaneous leishmaniasis in Somali region of Ethiopia

**DOI:** 10.1101/2024.10.15.617758

**Authors:** Adugna Abera, Pieter Monsieurs, Myrthe Pareyn, Dereje Beyene, Geremew Tasew, Allison Aroni-Soto, Mahlet Belachew, Desalegn Geleta, Bethlehem Adinew, Bokretsion Gidey, Henok Tadesse, Atsbeha Gebreegziabxier, Rajiha Abubeker, Abraham Ali, Ketema Tafess, Tobias F. Rinke de Wit, Johan van Griensven, Jean-Claude Dujardin, Dawit Wolday, Malgorzata Anna Domagalska

## Abstract

We sequenced *Leishmania tropica* genomes directly from eight human skin samples collected in a newly emerging focus of cutaneous leishmaniasis in Somali region, Ethiopia. We found a new variant with unique genomic signatures of drug resistance. Genomic homogeneity in the sampled population demonstrates the occurrence of a *L. tropica* outbreak.

## Text

Since the successful running of the Kala-Azar elimination program in the Indian sub-continent, the hotspot of worldwide leishmaniasis has moved to East Africa. Among the different affected countries, Ethiopia deserves particular attention, given the heterogeneous eco-epidemiology of leishmaniasis, its clinical polymorphism and the complex taxonomy of the parasites. The disease is endemic in different biotopes from low-to highlands and transmission involves different hosts and vectors. Four major clinical forms are encountered: visceral leishmaniasis (VL) and localized-, diffuse-and muco-cutaneous leishmaniasis (LCL, DCL and MCL, respectively). *L. donovani* (VL and occasionally CL) and *L. aethiopica* (LCL, DCL and MCL), are the most reported species, and *L. tropica* (CL) was signaled once in humans (1); in addition, several hybrids have been observed in isolates: *L. donovani/L. aethiopica* and *L. aethiopica/L. tropica*

The epidemiology of the disease is quite dynamic: it is affected by human migration and displacement due to famine and regular conflicts in the country and by environmental changes. We previously highlighted the need for genomic surveillance of leishmaniasis, using highly sensitive, resolutive and untargeted whole genome sequencing (WGS) methods (4). This recommendation is justified by several reasons: (i) since the discovery of hybridization and genetic introgression (5), robust species identification should theoretically be based on multi-genic approaches covering several regions of the genome (and not only single gene approaches); (ii) WGS is needed to assess the genetic similarity among parasites sampled from different patients, hereby confirming the outbreak nature of a focus and (iii) WGS can be used to find signatures of drug resistance and guide patient management.

In August 2023, an outbreak of CL was detected among immunologically naive militia that were recently deployed in the Eastern Somali region of Ethiopia, in an area without prior history of CL cases, yet sporadic cases of VL were reported. The clinical presentation of CL was not comparable to what is typically observed in Ethiopia and neighboring countries. In first instance the CL pathogen was identified by Hsp70 amplicon sequencing as *L. tropica* (3).

We aimed here to undertake a more in-depth molecular characterization of *L. tropica* samples collected in the Somali focus. We applied a method of direct genome sequencing of *Leishmania* in host tissues (SureSelect sequencing [SuSL-seq]; Agilent Technologies, https://www.agilent.com), not requiring parasite isolation and cultivation (6). The method was so far validated for *L. donovani* in bone marrow (6) and blood samples (7) and was here applied for the first time to skin samples from CL patients.

Eight of the Somali CL samples were submitted to SuSL, using a capture panel of probes designed for *Leishmania aethiopica* genome; noteworthy, this library should work very well with phylogenetically related species like *L. tropica*. In a first step, species identification of the 8 samples was done by a phylogenetic analysis, using as references the published genomes of *L. aethiopica, L. tropica, L. major, L. donovani* and inter-species hybrids (appendix). As shown in Figure S2, the eight Somali samples (yellow arrow) clearly branch in the *L. tropica* cluster and they are genetically very different from *L. aethiopica, L. major, L. donovani* and inter-species hybrids. In a second step, we focused on *L. tropica* genomes only (Fig.1A). This provided three major insights: (i) the eight Somali parasites constitute a *L. tropica* variant not reported so far in analyzed genomes, (ii) they branch between genotypes reported so far from Israel/Jordan and other Middle East variants, (iii) they cluster together and are genetically very homogeneous (on average 122 SNPs between samples), which confirms their outbreak origin. Finally, we found homozygous missense mutations or frameshifts in 14 genes reported to be involved in drug resistance, especially antimony (Fig.1B). This signature was never reported before in any *L. tropica* genome, which confirms the unique character of Somali samples. Further work is required to understand the clinical impact of this discovery.

**Fig. 1A.**
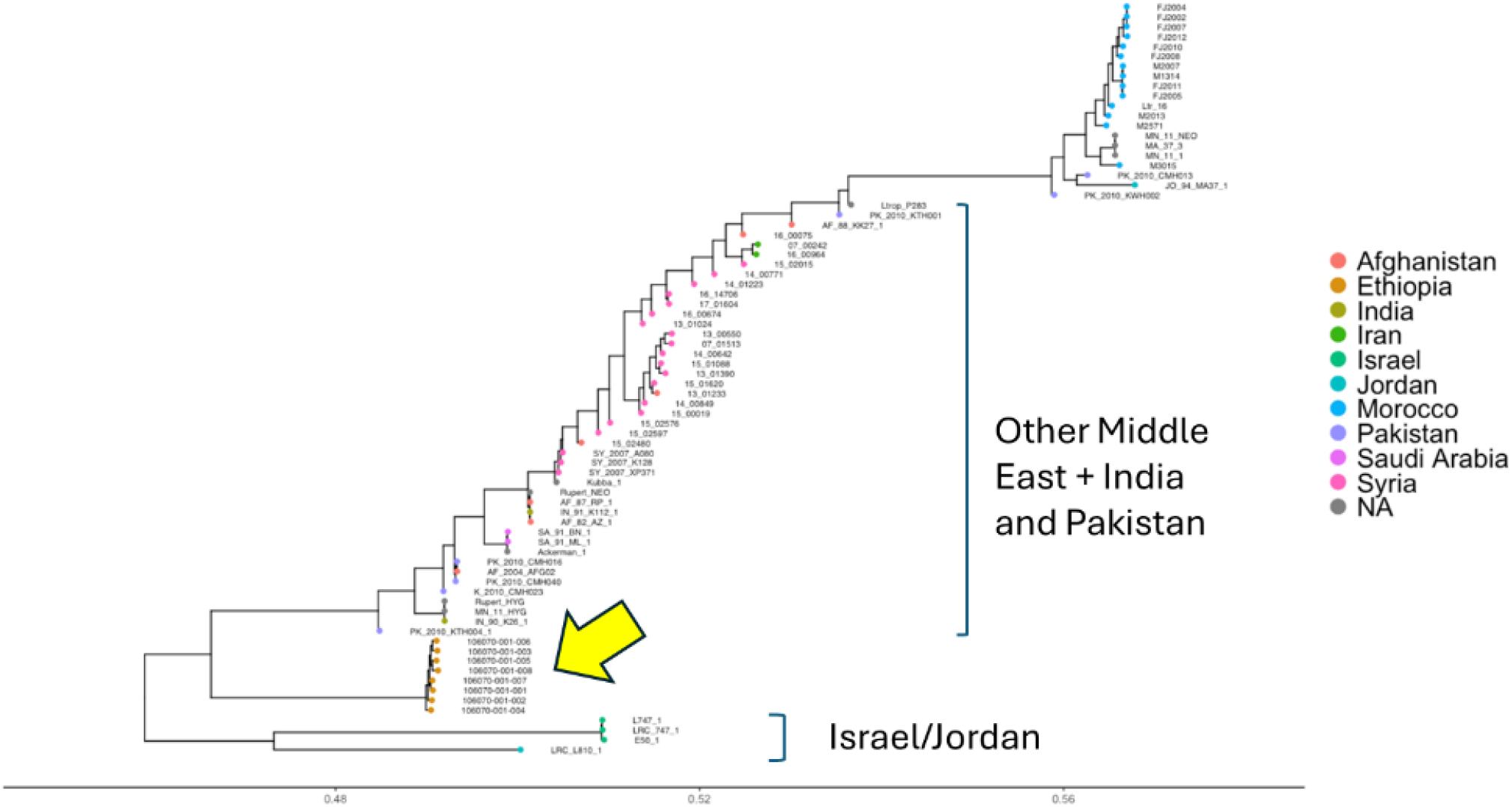
Rooted phylogenetic tree of all publicly available *L. tropica* genomes, generated with RAxML with the GTR+G substitution model, and showing the close clustering of the eight samples presented in this work (indicated by a yellow arrow). Important bootstrap values are indicated on the branches. The *L. infantum* JPCM5 reference genome was included as an outgroup but is omitted from the figure. Scale bars indicate number of single-nucleotide polymorphism differences. Color code represents the country of origin of the strain.

**Fig. 1B.**
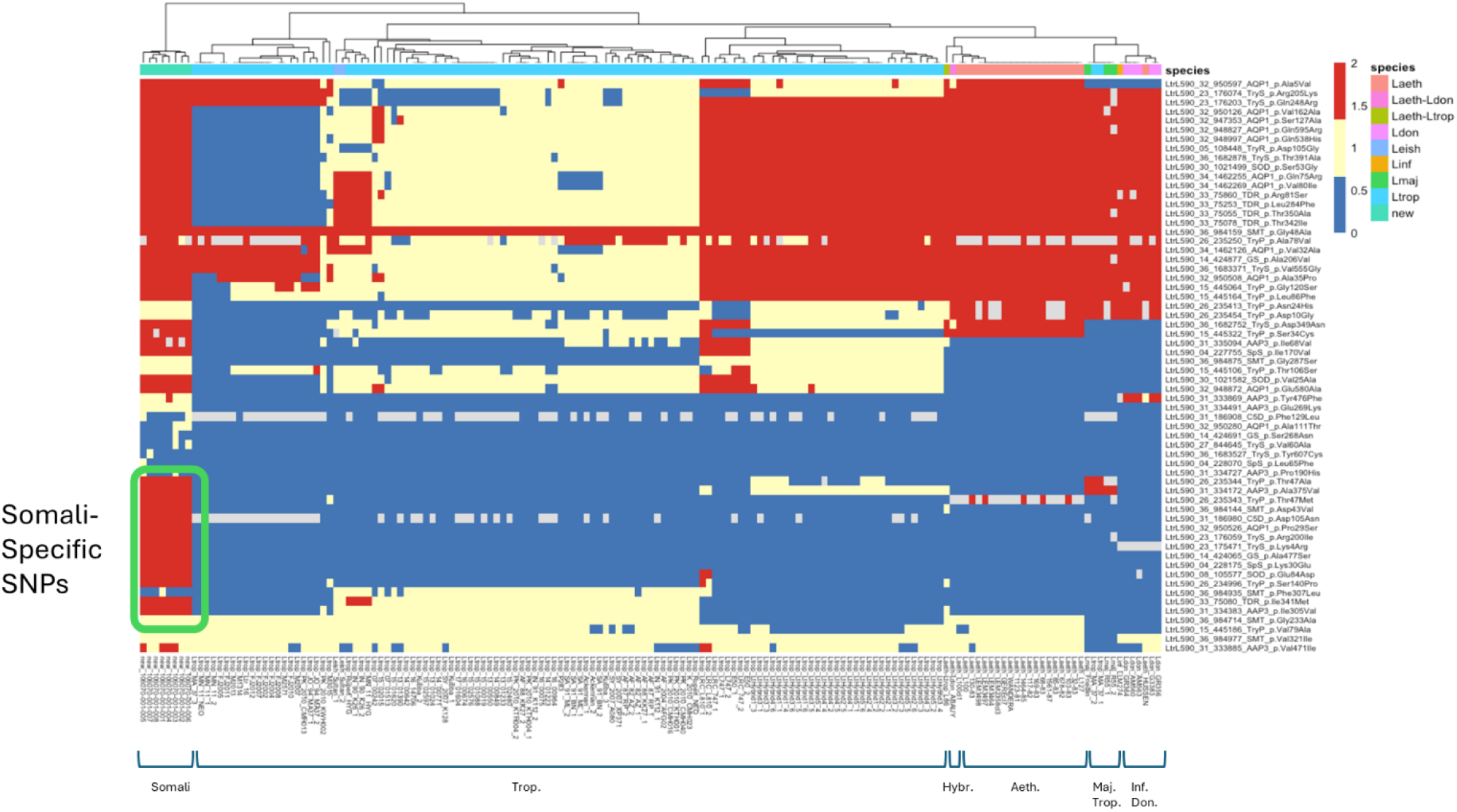
Heatmap showing the distribution of single nucleotide polymorphisms (SNPs) in genes reported to be associated with drug-resistant phenotypes (see appendix). The color scheme represents different SNP categories: blue indicates the absence of SNPs, orange indicates heterozygous SNPs, and red indicates homozygous SNPs. The naming convention for SNPs follows the format of the gene of interest, position in the genome, type of mutation, and its effect on the corresponding protein. Trop., *L. tropica*; Hybr., hybrids (between *L. aethiopica* and *L. tropica* or *L. donovani*; Aeth., *L. aethiopica*; Maj., *L. major*; Inf., *L. infantum*; Don., *L. donovani*

*L. tropica* is essentially endemic in Morocco, Turkey, Syria, Israel, Iraq, Azerbaidjan, Iran, Uzbekistan, Afghanistan, Pakistan, India (8). This broad distribution likely results from the anthroponotic nature of *L. tropica* transmission and the old communication axes in many of these countries, like the Silk Road. In some regions, sporadic cases are reported and the disease is thought to be zoonotic, with possible animal reservoirs including hyraxes, bats or wild rodents (9). The high genomic homogeneity in the population here sampled demonstrates the occurrence of a *L. tropica* outbreak in the Somali region of Ethiopia. Our study does not allow yet to trace the origin of the outbreak, being a primary human case from which the parasite population would have spread or being an animal reservoir. It highlights the risk of further expansion of the parasites from the human cases in this new focus and the need for genomic surveillance in humans and animals, also in neighbor countries, like Kenya where *L. tropica* was recently reported (10).

Genomic sequence reads of the parasites from the 8 samples have been submitted to the Sequencing Read Archive of NCBI (https://www.ncbi.nlm.nih.gov/sra). Bioproject: PRJNA1172382

This study was financially supported by (i) the Belgian Directorate-General for Development Cooperation Framework Agreement 5 (FA5) Ethiopia program, awarded to JvG and GT, (ii) the Dioraphte foundation (Spatial-CL, project number: CFP-RD2020 20020401), (iii) EpiGen-Ethiopia (Project ID#101103188, funded through the Global Health EDCTP3 program - European Union) and (iv) the Flemish Ministry of Science and Innovation, support to MAD.

## About the author

Dr. Abera is a senior researcher at the Ethiopian Public Health Institute since 2016. He focuses on molecular epidemiology, drug resistance, diagnostics, and metagenomics research applied to malaria, NTDs, and arboviral diseases in Ethiopia.

