## appendix for "Genomic characterization of a unique population of *Leishmania tropica* in a new emerging focus of cutaneous leishmaniasis in Somali region of Ethiopia"

**Appendix for submission**

This document contains:

A1. Clinical procedures and data

A2. Laboratory procedures

A3. Bioinformatic procedures

A4. Additional results

A5. References

**A1. Clinical procedures and data**, including summary of clinical and epidemiological data from patients (Table S1) and a map showing their geographical origin (Fig.S1); more details can be found elsewhere (1).

Eight male members of the local militia who participated in the border conflict between the Somali and Afar regions were studied. Each referral facility had a dermatologist who examined suspected cases of cutaneous leishmaniasis (CL). The lesions were categorized as localized cutaneous leishmaniasis (LCL), or mucocutaneous leishmaniasis (MCL) (Table S1). Before starting treatment, fine needle aspirate (FNA) and skin scraping samples were taken from each case by experienced health professionals for Giemsa smear preparation. The Giemsa-stained results tested positive for leishmania amastigotes.

**Table S1.** Clinical data of the 8 patients here studied.

*JUSYCSH (Jigjiga University Sheik Hassen Yabare Comprehensive Specialized Hospital); DHC (Duunyar Health Center); SPH (Sitti Primary Hospital)*

Except for the two patients diagnosed at Duunyar Health Center, all six patients were treated with systemic intramuscular sodium stibogluconate (SSG) and were followed for 28 days. All six patients responded well to the SSG treatment, and no relapse cases have been reported from the treating facilities.

| **Specimen ID** | **Sex** | **Age** | **Facility Name** | **Site of lesion** | **Number of leisions** | **Types of CL** | **Duration since onset of lesion in months** | **Giemsa stained result** |
|---|---|---|---|---|---|---|---|---|
| WH08 | Male | 40 | JUSYCSH | Face | 12 | LCL | 6 | Positive |
| WH09a | Male | 20 | JUSYCSH | Face, Nose, Lips | 13 | MCL | 8 | Positive |
| WH11 | Male | 22 | JUSYCSH | Face | 8 | LCL | 6 | Positive |
| WH12 | Male | 40 | JUSYCSH | Face, Nose, Lips | 14 | MCL | 7 | Positive |
| S04 | Male | 29 | DHC | legs | 3 | LCL | 6 | Positive |
| S06 | Male | 42 | DHC | Face, Nose, Lips | 18 | MCL | 6 | Positive |
| S12 | Male | 36 | SPH | Face, Nose, Lips | 11 | MCL | 5 | Positive |
| WH15 | Male | 21 | JUSYCSH | Face | 10 | LCL | 6 | Positive |

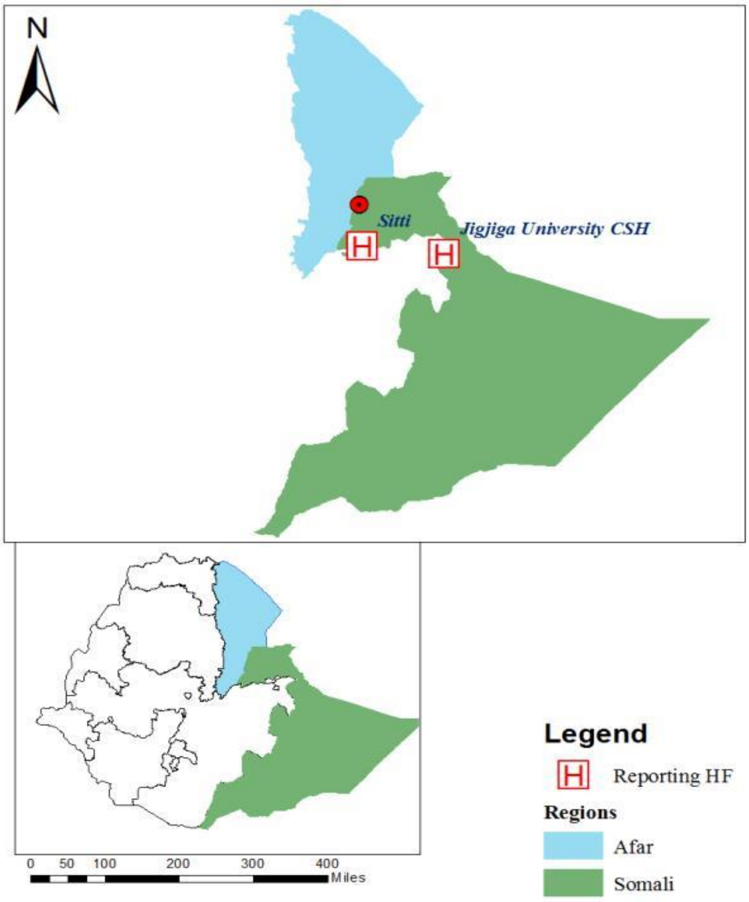


**Fig S1**: Study site indicating the locations of the healthcare facilities for the eight study subjects in the Somali region of Ethiopia. The map of Ethiopia shows the Afar region in blue and the Somali region in green. Two cases (S04 and S06) were diagnosed at Duunyar Health Center, located at the border of the Afar and Somali regions, where the outbreak occurred (indicated by a red dot). One case (S12) was diagnosed at Sitti Primary Hospital, and five cases (WH08, WH09a, WH11, WH12, and WH15) were reported at Jigjiga University Sheik Hassen Yabare Comprehensive Specialized Hospital, marked by red "H".

### A2. Laboratory procedures

We received 37 samples, selected 8 (Table S2), using as criteria a minimum of 20 ng of input gDNA, and the % of Leishmania DNA at least 0.006% measured by qPCR as described elsewhere (2).

**Table S2.** Technical features of the samples here studied

| Samples | Total quantity of DNA (ng) | % Leishmania DNA | Enrichment |
|---|---|---|---|
| WH08 | 6860 | 0,016 | no |
| WH09a | 1981 | 0,024 | no |
| WH11 | 1071 | 0,053 | no |
| WH12 | 1089 | 0,014 | no |
| S04 | 51 | 0,152 | Ampure Beads |
| S06 | 117 | 0,438 | Ampure Beads |
| S12 | 22 | 0,081 | Ampure Beads |
| WH15 | 165 | 0,014 | Ampure Beads |

A custom capture panel of probes with a size of 26.55 Mbp was designed for *Leishmania aethiopica* genome, using the reference genome sequence of the L146 strain and synthesized by Agilent Technologies. Our *L. aethiopica* array should be suitable for analysing samples with both *L. aethiopica* and *L. tropica* infections. Indeed, *L. tropica* and *L. aethiopica* are phylogenetically highly related to each other (same species complex). Furthermore, we successfully applied an array developed using mostly *L. infantum* genome for capture and sequencing of *L. donovani* genome (also in a same species complex) (2).

The library preparation was done following SureSelect XT HS Target Enrichment System for Illumina Multiplexed Sequencing Platforms protocol (Agilent Technologies, Santa Clara, USA). First, samples with the starting concentration of the DNA between 0.7–3.9 ng/µl), were concentrated via purification on Ampure Beads and the elution in 10µl of low TE. In brief, quantity of 10-200 ng of input gDNA was fragmented using SureSelect Enzymatic Fragmentation kit (Agilent technologies, Santa Clara, USA). Subsequently, adaptor-ligated libraries were prepared using 8,10 or 12 number of cycles in pre-capture PCR, depending on the amount of input DNA. Libraries were hybridized with the custom probes at a dilution 1:10, and captured with Dynabeads MyOne Streptavidin T1 magnetic beads (Thermo Fisher Scientific, Waltham, USA). After washing steps, the DNA captured by streptavidin beads was amplified by PCR, and purified with AMPure XP beads. The quantity and quality of the libraries were assessed by TapeStation using High Sensitivity D1000 ScreenTape (Agilent Technologies, Santa Clara, USA). Sequencing was done in Genomescan (Leiden, the Netherlands) using Illumina NovaSeq™ 6000 platform, 150 bp paired-end reads.

**A3. Bioinformatic procedures**

Clinical samples for this study underwent PCR-free whole-genome sequencing utilizing the Illumina NovaSeq platform, producing 2x150 bp paired-end reads. Additionally, we incorporated previously published sequencing datasets, as available in the NCBI's Sequence Read Archive (SRA). This included *L. tropica* genomes from BioProjects PRJEB6281, PRJEB45563, and PRJNA978932, as well as genomes of *L. aethiopica* and other *Leishmania* species from BioProject PRJNA924694. The SRAtoolkit software was employed for downloading these publicly accessible sequencing data. Reads were aligned to the *L. tropica* L590 reference genome (TriTrypDB version 58) using BWA (v0.7.17) with a seed length of 50 (3). We selected only properly paired reads with a mapping quality score exceeding 30, processed using SAMtools (4). Duplicate reads were eliminated using the RemoveDuplicates feature in Picard software (v2.22.4). SNP calling followed the Genome Analysis ToolKit (GATK) best practices (v4.1.4.1). The procedure included: 1) Utilizing GATK

HaplotypeCaller to generate GVCF files for each sample. 2) Combining these GVCF files with the GATK CombineGVCF command. 3) Performing genotyping via GATK GenotypeGVCF. 4) Filtering SNPs and indels as per GATK's "best practices" using SelectVariants and VariantFiltration commands. For constructing phylogenetic trees, biallelic SNPs from VCF files were selected using BCFtools and converted to Phylip format with the vcf2phylip.py script. RAxML was employed with the GTR+G substitution model, performing 100 bootstrap replicates. *L. infantum* JPCM5 was used as an outgroup. The resulting trees were visualized using ggtree for rooted phylogenetic trees and SplitsTree for unrooted phylogenetic networks.

**A4. Additional results**

Species identification was made by aligning -using BWA (3)- the sequencing reads from all eight samples to artificial concatenated genome consisting of 1) the human genome, 2) the *L. tropica* genome L590 as downloaded from TriTrypDB version 68, 3) the *L. aethiopica* L147 reference genome as downloaded from TriTrypDB version 68. Using this competitive mapping approach, the BWA algorithm decides to which genome each sequencing reads maps with the highest quality. Using SAMtools (4), only those sequencing reads uniquely mapping to the artificial reference genome are selected. The relative percentage of reads mapping to either the *L. tropica* L590 or the *L. aethiopica* L147 reference genome is used to derive the species and exclude the potential hybrid nature of the strains.

**Fig.S2.** Phylogenetic network indicating the phylogenetic relation between a subset of different Leishmania genomes (strains belonging to *L. tropica*, *L. aethiopica*, *L. major* and *L. donovani*) The quality-filtered SNP VCF files were transformed into FASTA format using the vcf2fasta.py script (accessible at github.com/FreBio/mytools/blob/master/vcf2fasta.py). To explore the phylogenetic relationships among the genomes, a phylogenetic network was generated with SplitsTree version 4.19.0 (5) based on concatenated bi-allelic SNPs. The eight newly sequenced genomes group together with the other *L. tropica* genomes. Apart from a previously identified hybrid between *L. aethiopica* and *L. donovani* (Abauy strain), we also found a potential previously unidentified hybrid based on its position in this phylogenetic network (Sultan_1)

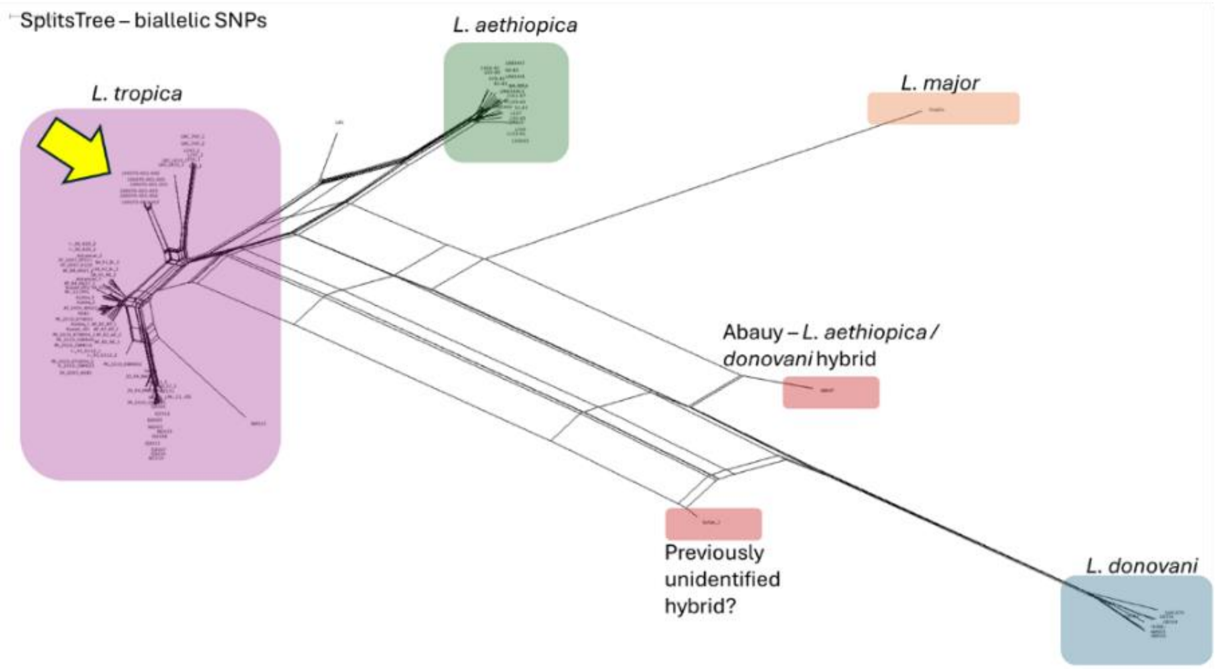

For the exploration of possible genomic signatures of drug resistance, we selected 11 loci that were previously shown to be involved in *L. tropica* resistance to antimonials, we complemented these with 4 loci reported to be involved in resistance to miltefosine and amphotericin B in other species. Relevant regions (indicated below) were subset using BCFtools (6). Only SNPs having an effect at the protein level (missense and non-sense mutations) were retained. Visualization of the heatmap (Fig.1B) was performed using the pheatmap function in R. ***For antimonials**:* Glutathione synthetase (GS)(7); Spermidine synthetase (SpS)(7); Thiol-dependant reductase (TDR)(7); Mitochondrial superoxide dismutase (SODA)(8); Glycosomal superoxide dismutase (SODB)(8); Tryparedoxin peroxidase (TryP)(9) ; Trypanothione reductase (TryR) (9); Aquaglyceroporine 1 (AQP1)(10); ABC transporter (MRPA)(10); Leishmania-activated C kinase gene (LACK1)(11); Amino acid permease (AAP3)(12). ***For miltefosine***; Miltefosine transporter (LdMT)(13); Beta-subunit of LdRos3(13). ***For Amphotericine B***: Sterol C5-desaturase (C5D)(14); Sterol C24-methyltransferase (SMT)(14)
